# Propofol modulates functional connectivity signatures of sustained attention

**DOI:** 10.1101/2021.10.15.464605

**Authors:** Taylor Chamberlain, Monica D. Rosenberg

## Abstract

Sustained attention is a critical cognitive function reflected in an individual’s whole-brain pattern of fMRI functional connectivity. However sustained attention is not a purely static trait. Rather, attention waxes and wanes over time. Do functional brain networks that underlie individual differences in sustained attention also underlie changes in attentional state? To investigate, we replicate the finding that a validated connectome-based model of individual differences in sustained attention tracks pharmacologically induced changes in attentional state. Specifically, preregistered analyses revealed that participants exhibited functional connectivity signatures of stronger attention when awake than when under deep sedation with the anesthetic agent propofol. Furthermore, this effect was relatively *specific* to the predefined sustained attention networks: propofol administration modulated strength of the sustained attention networks more than it modulated strength of canonical resting-state networks and a network defined to predict fluid intelligence, and the functional connections most affected by propofol sedation overlapped with the sustained attention networks. Thus, propofol modulates functional connectivity signatures of sustained attention within individuals. More broadly these findings underscore the utility of pharmacological intervention in testing both the generalizability and specificity of network-based models of cognitive function.

## Introduction

Attention fluctuates within individuals both due to natural changes in arousal, such as dozing off during a long lecture, but also due to pharmacological intervention, such as in the use of methylphenidate in the treatment of attention deficit hyperactivity disorder (ADHD). Pharmacologically induced differences in attentional state have measurable real-world consequences; children who are prescribed methylphenidate for ADHD showing significant improvement in academic performance metrics such as math accuracy and reading speed (Kortekaas-Rijlaarsdam et al., 2018).

To what extent are pharmacologically induced changes in cognitive and attentional state reflected in patterns of whole-brain functional connectivity? Work suggests functional connectivity is dominated by stable individual traits, with relatively little contribution of state-related variance to an individual’s functional connectome. For instance, Gratton et al. contrasted subject-dependent, task-dependent and session-dependent variation in the functional connectome and found that the vast majority of variation could be attributed to subject-dependent effects (Gratton et al., 2018).

At the same time, other work has provided evidence that functional connectivity does, in fact, vary meaningfully with changes in mental states. For instance, recent work found an increase in network strength of a functional connectivity-based model of mind-wandering over the course of four runs of rest scans, in tandem with a decrease in ratings of thoughts related to the external world (Kucyi et al., 2021). The sustained attention connectome-based predictive model (CPM), a model of individual differences in sustained attention, has also been shown to be sensitive to within-subject attention changes (Rosenberg, Finn, et al., 2016). The model consists of two functional networks, both defined in a data driven manner, which predict better and worse attention, respectively. The model has been validated in its prediction of individual differences in attention function across multiple independent datasets, and shows sensitivity methylphenidate, such that individuals given a single dose before functional MRI show functional connectivity signatures of better sustained attention (Rosenberg, Zhang, et al., 2016). Furthermore, individuals under deep sedation with propofol and light anesthesia with sevoflurane showed functional connectivity signatures of worse sustained attention compared to when they were resting while awake (Rosenberg et al., 2020).

Here, with a preregistered replication in an independent fMRI dataset, we add to this growing body of evidence by examining the effect of propofol on the sustained attention CPM—as well as the specificity of this effect—during both rest and a naturalistic task. Additionally, we test the hypothesis that connectome-based models do not always capture state-like and trait-like variability in the behavior they were defined to predict. Instead, networks defined to predict more trait-like abilities, such as fluid intelligence, may be less sensitive to within-subject cognitive and attentional state changes. Finally, we investigate the effect of propofol on functional connectome patterns more broadly, characterizing the degree to which propofol administration increases or decreases connectome similarity between individuals and across task states.

## Methods

### Dataset

We performed secondary analyses of data available on Openneuro.org (https://openneuro.org/datasets/ds003171) (Kandeepan et al., 2020; Naci et al., 2018). In this dataset, functional MRI data were acquired while participants rested and listened to a 5:12-minute audio clip from the film *Taken* at four different levels of sedation with the anesthetic agent propofol.

### Participants

Seventeen healthy, right-handed, native English speakers (4 women; mean age 24 years, SD = 5) participated in the original study, which was approved by the Health Sciences Research Ethics Board and Psychology Research Ethics Board of Western University (REB #104755). All participants completed an MRI and propofol safety screening questionnaire provided by both the attending MR technician and anesthesiologist, provided informed consent, and were paid for their participation. Secondary analysis of these data was approved by the University of Chicago Institutional Review Board. Scans for which more than 50% of frames were censored for head motion (see *Functional MRI data preprocessi*ng) were excluded from analyses. Of the seventeen participants, ten had *awake* and *deep sedation* resting-state scans which passed motion exclusion, and ten partially overlapping participants had *awake* and *deep sedation* narrative-listening scans. All analyses were performed on these two sets of ten participants, with the exception of the *Functional connectivity similarity analyses* (see below).

### Task protocol

Functional MRI scans were acquired during four different levels of sedation: *awake, mild, deep*, and *recovery*. During each level of sedation, rest and narrative-listening scans were acquired (8 and 5 minutes respectively). During the rest scan, participants were instructed to relax with their eyes closed without falling asleep. During the narrative listening scan, participants listened to an audio excerpt from the movie *Taken*. In this emotionally evocative clip, listeners hear a teenage girl being kidnapped while speaking to her father on the phone. During each of the four sedation conditions, the narrative scan preceded the rest scan.

### Propofol administration and sedation assessment

Before fMRI data acquisition for each of the four levels of sedation, two anesthesiologists and one anesthesia nurse evaluated volunteers’ Ramsay level, which classifies a person’s level of sedation on a scale from one (severe agitation) to six (deep coma). Scanning for each session began once the three anesthesia assessors agreed on the participant’s sedation level. During the *awake* session, no propofol was administered. Propofol infusion began prior to the *mild* session, and the *mild* session commenced once participants reached Ramsay 3 level of sedation, in which participants’ response to verbal communication slowed. Prior to the *deep* session, propofol target effect-site concentration was increased until participants reached a Ramsay 5 level of sedation, in which participants stopped responding to verbal commands. Following the *deep* sedation scan, propofol infusion was discontinued and once a Ramsay 2 level of sedation was achieved, the *recovery* session commenced. At Ramsay level 2, participants exhibited quick responses to verbal commands. For detailed descriptions of propofol administration protocol see (Naci et al., 2018).

### Functional MRI data acquisition

Participants wore noise-canceling headphones, and volume was adjusted to each participant’s level of comfort (Naci et al., 2018). MRI data were acquired with a 3-Tesla Siemens Tim Trio scanner (32-channel coil). Functional images were collected with the following parameters: voxel size = 3 × 3 × 3 mm^3^, inter-slice gap of 25%, TR = 2000 ms, TE = 30 ms, matrix size = 64 × 64, FA = 75°. narrative scans and resting-state scans had 155 and 256 volumes respectively. Anatomical images were acquired as well, with a T1-weighted 3D MPRAGE sequence (32 channel coil, voxel size: 1 × 1 × 1 mm^3^, TE = 4.25 ms, matrix size = 240 × 256 × 192, FA = 9°).

### Preregistration

Primary hypotheses, planned tests, and fMRI preprocessing steps were preregistered on the Open Science Framework prior to data analysis (https://osf.io/ndh3v/registrations). Hypotheses and tests described in the *Network strength calculation* and *Network strength as a function of sedation level* sections of the *Methods* were preregistered. Follow-up analyses described in the *Specificity of propofol effects, Propofol network identification*, and *Representational similarity analyses* sections were not preregistered.

### Functional MRI data preprocessing

Functional MRI preprocessing steps (with two minor changes, detailed below) were preregistered prior to data analysis. AFNI was used to preprocess functional MRI data. First, three volumes were removed from each run, followed by despiking, and head motion correction. Then, functional images were aligned to the skull-stripped anatomical image with a linear transformation and then to the MNI atlas via nonlinear warping. Covariates of no interest were regressed from the data, including a 24-parameter head motion model (6 motion parameters, 6 temporal derivatives, and their squares) and mean signal from subject-specific eroded white matter and ventricle masks and the whole brain. Because head motion in this sample was relatively high (mean frame-to-frame displacement before participant, run, or frame exclusion = 0.153 mm; Supplementary Figure 1), the final preprocessing pipeline deviated from the preregistered pipeline in two ways: the addition of censoring of high-motion volumes and the removal of band-pass filtering. Volumes in which more than 10% of voxels were outliers and volumes for which the Euclidean norm of the head motion parameter derivatives exceeded 0.25 were censored from the time-series. Voxel-wise BOLD signal time-courses were averaged within regions of interest using a 268-node whole-brain parcellation (Shen et al., 2013).

### Network strength calculation

Functional network nodes were defined with the 268-node functionally defined whole-brain Shen atlas (Shen et al., 2013). We conducted our primary replication analysis in two ways: first, including all 268 nodes, and second, including 238 nodes, dropping any node (30 in total) that was missing in any scan (see Supplementary Figure 2 for the latter analysis). Functional connectivity, defined as the Fisher *z*-transformed Pearson correlation between the fMRI signal time-courses of pairs of atlas parcels, was calculated for each fMRI run separately.

### Network strength as a function of sedation level

To characterize the degree to which volunteers expressed functional connectivity signatures of sustained attention defined in previous work, sustained attention network strength was measured in each functional connectivity matrix. This was performed with the high- and low-attention network masks, available at [https://github.com/monicadrosenberg/Rosenberg_PNAS2020], which comprise the predefined sustained attention connectome-based predictive model (CPM). These masks consist of 268 × 268 binary matrices where a value of one indicates a functional connection, or edge, in the mask. We applied each mask to the functional connectivity matrix, then averaged the values in each network for each functional connectome separately, yielding high- and low-attention strength values. Prior work has demonstrated that attention network strength tracks both interindividual and intraindividual differences in sustained attention, where higher high-attention network and lower low-attention network scores are associated with better sustained attention function (Rosenberg et al., 2020; Rosenberg, Finn, et al., 2016). This analysis resulted in our main variables of interest: eight separate high-attention and low-attention network strength values for each participant with complete data (two tasks [rest, narrative listening] x four sedation conditions [*awake, mild sedation, deep sedation, recovery*]).

To directly replicate previous work showing that propofol sedation significantly modulates sustained network strength (Rosenberg et al., 2020), we performed a paired *t*-tests comparing high-attention network strength in the *awake* and *deep sedation* conditions and low-attention network strength in the *awake* and *deep sedation* conditions during rest. To test if this effect generalized to a different task state, we repeated both tests with narrative-listening data.

Although the *t*-tests directly replicate the analysis of Rosenberg et al. (2020), they have the disadvantage of excluding two of the four levels of sedation (*mild, recovery*), and fail to test for possible interactions between the effect of task and sedation. To examine the effect of all four levels of sedation and task manipulations simultaneously, we assessed the main effects of sedation level and task and their interaction with two mixed-effects models using the lme4 package in R (Bates et al., 2015), one for normalized high-attention network strength and one for normalized low-attention network strength. Sedation level and task were included as fixed effects, and participants were included as random effects. Including random slopes for participants with respect to the effect of sedation prevented model convergence, and including random slopes for participants with respect to the effect of task did not improve (i.e., decrease) the model’s AIC. Thus, random slopes were not included in the models.

### Specificity of propofol effects

We tested whether any effects of propofol administration were specific to networks that predict sustained attention in three ways. First, we examined the effect of sedation on a connectome-based model defined to predict another central cognitive measure, fluid intelligence (gF) (Greene et al., 2018). This fluid intelligence CPM was defined using fMRI data from the Human Connectome Project sample (collected while participants performed an *n*-back working memory task) to predict individual differences in performance on a 24-item version of the Penn Progressive Matrices test. Fluid intelligence has been shown to be relatively stable across the lifespan and is not thought to vary from one moment to the next (Kazlauskaite & Lynn, 2002; Schaie et al., 2004). Consequently, networks that predict fluid intelligence may not vary with short-term state changes, akin to the kind induced by propofol, to the same degree as do networks defined to predict attention. Thus, we predicted that strength in the fluid intelligence networks would be less affected by the sedation manipulation than strength in the sustained attention networks. To test this hypothesis, we repeated the analysis described above to generate high- and low-fluid intelligence network strength values for every fMRI run and compared these values between sedation conditions.

Second, replicating previous work (Rosenberg et al., 2020), we calculated strength in canonical resting-state networks (defined in Finn et al., 2015), such as the default mode network and frontoparietal network, to compare the relative effect of sedation on these networks with the sustained attention network.

Third, we identified sets of functional connections that significantly differed between individuals’ *awake* and *deep* sedation scans, and asked whether these networks overlapped with the predefined sustained attention networks (see *Propofol network identification* and *Network overlap* sections below). We predicted that a network whose strength decreased with propofol administration would overlap with the network predicting better sustained attention, whereas a network whose strength increased with propofol administration would overlap with the network predicting worse sustained attention.

### Propofol network identification

Using the Network Based Statistic (NBS) Toolbox (RRID:SCR_002454; <u>sites.google.com/site/bctnet/comparison/nbs</u>), we identified functional networks that differed between the *awake* and *deep sedation* conditions. This procedure aids network detection by controlling for the large number of multiple comparisons necessary to test for differences in every edge in a functional connectome, by comparing the size of the fully connected networks that differ between two conditions of interest (here, *awake* and *deep sedation*) to the size of fully connected networks that differ between randomly assigned conditions (Zalesky et al., 2010).

First, we identified edges that were greater in the *deep sedation* or *awake* condition by performing paired, one-tailed *t*-tests, and retaining edges that fell above a predetermined significance threshold. Then we selected the largest fully connected network of edges from this group. Next permutation testing was performed by shuffling condition labels 5000 times and running a *t*-test at every permutation to generate 5000 sets of edges which differed between the random groups. The *p* value of the sedation network was calculated by computing (*n_fully_connected_random* + 1) / (*n_permutations* + 1), where *n_fully_connected_random* is the number of fully connected random network components the same size or larger than the observed fully connected network component, and *n_permutations* is 5000, the number of permutations. We classified the resulting networks as significant at *p* < 0.05.

We repeated this process for each task condition (rest and narrative listening), and for three different significance thresholds at the edge-selection step (*p* < 0.001, *p* < 0.01, and *p* < 0.05), resulting in three *Awake* and three *Deep Sedation* networks for each task.

### Network overlap

Considering the marked impacts of sedation on cognitive and attentional states, we predicted significant overlap between the high-attention network and the *Awake* propofol network, and significant overlap between the low-attention network and the *Deep Sedation* propofol network. Furthermore, we expected less overlap between the propofol networks and the fluid intelligence networks, since the fluid intelligence networks may be more reflective of trait-like differences in functional connectomes rather than the temporary state changes in the connectome caused by propofol sedation.

To test these hypotheses, we examined the overlap between the propofol networks and the sustained attention and fluid intelligence networks. For each pair of networks, we counted the number of edges shared between the networks. The significance of the overlap was determined with the hypergeometric cumulative density function, which provides the probability of drawing up to *x* of *K* possible items in *n* drawings without replacement from a population with *M* items. This was implemented in MATLAB as *p* = 1 – *hygecdf*(*x, M, K, n*), where *x* is the number of overlapping edges, *K* is the number of connections in the given CPM network, *n* is the number of connections in the given propofol network, and *M* is the total possible number of edges in the matrix (35,778, all possible functional connections given 268 nodes). We repeated this process for all three versions of the propofol networks (thresholded at *p* < .05, *p* < .01, and *p* < .001 respectively) and both tasks (rest, narrative). To determine which CPM (sustained attention or fluid intelligence) had greater overlap with the propofol networks, permutation testing was performed. We compared the difference in percent overlap between the networks by shuffling the propofol network 1000 times and using the amount of overlap with the randomly generated network as a null distribution. Again, this was repeated for all propofol networks and both task conditions.

### Functional connectivity similarity analyses

In addition to examining effects of task state and sedation level on functional connectivity signatures of sustained attention, we were interested in understanding the effects of the drug and task manipulations on functional connectivity patterns more broadly. That is, how does sedation and task state affect the similarity of functional connectivity patterns across individuals? Do individuals show more similar connectivity patterns when awake or when under sedation? One possibility is that sedation *increases* similarity across individuals by reducing ongoing cognitive, attentional, and perceptual processes that may otherwise differ between people and drive unique patterns of functional brain organization. Another is that sedation *decreases* similarity between individuals by muting these processes, revealing idiosyncratic underlying patterns of functional brain organization. In the same vein, do individuals show more similar connectivity patterns when listening to the same story or resting? We might expect that participants listening to the same story would exhibit more similar patterns of functional connectivity because they are engaged in similar auditory and cognitive processing.

To investigate how propofol sedation affects between-subject connectome similarity, participants’ fMRI time series were limited to 73 TRs per run to match the number of TRs per condition. All nodes missing in any scan were excluded from analysis (30 in total). Next, for both task conditions, we excluded participants who did not pass our motion threshold (< 50% of frames censored for motion) for the *awake* and *deep sedation* conditions, resulting in 10 (rest), and 10 (narrative listening) participants, respectively (7 participants included in both conditions). We then calculated the Spearman correlation of each participant’s functional connectivity pattern for the *awake* resting-state scan with that of all other participants. We averaged the Fisher-*z* transforms of these values to get a single similarity value per participant. We repeated this process for the *deep* condition and for the narrative listening task, yielding four lists of subject-level similarity values. Finally, similarity while participants were *awake* was compared to similarity under *deep sedation* using two paired *t*-tests, one for each task condition.

To investigate how the task manipulation affects between-subject similarity, we again limited participants’ time series to 73 TRs, excluding all nodes missing in any scan. Next, for each of the four sedation conditions, we excluded participants who did not pass our motion threshold for both task conditions, resulting in 16 (*awake*), 13 (*mild*), 8 (*deep*), and 15 (*recovery*) participants respectively (6 participants in all conditions). We calculated the Spearman correlation of each participant’s functional connectivity pattern during the *awake* resting-state scan with that of all other participants. We averaged the Fisher-*z* transforms these values to get a single similarity value per participant. We repeated this process for all four sedation conditions and for the narrative-listening task, resulting in eight lists of subject-level similarity values. Finally, similarity during resting-state scans was compared to similarity during narrative listening using four paired *t*-tests, one for each sedation level.

## Results

### Propofol decreases functional connectivity signatures of sustained attention

As predicted, during rest, high-attention network strength was higher (*t*_9_ = 3.22, *p* = 0.010) and low-attention network strength was lower (*t*_9_ = -4.09, *p* = 0.003) in the *awake* than the *deep sedation* condition. This pattern of results replicated during narrative listening (high-attention: *t*_9_ = 2.50, *p* = 0.034; low-attention: t_9_ = -3.83, *p* = 0.004).

Also as predicted, mixed effect models of propofol’s effect on sustained attention network strength revealed a main effect of sedation condition wherein less sedation was associated with functional connectivity signatures of stronger attention (high-attention: *b* = -0.75, SE = 0.16, F[3,93.5] = 8.01, *p* = 8.26×10^−5^; low-attention: *b* = 0.84, SE = 0.15, F[3,92.4] = 13.24, *p* = 2.91×10^−7^; **Fig. 1**). There was no main effect of task (i.e., rest vs. narrative listening) or interaction of sedation condition and task on high-attention or low-attention network strength.

**Figure 1.**
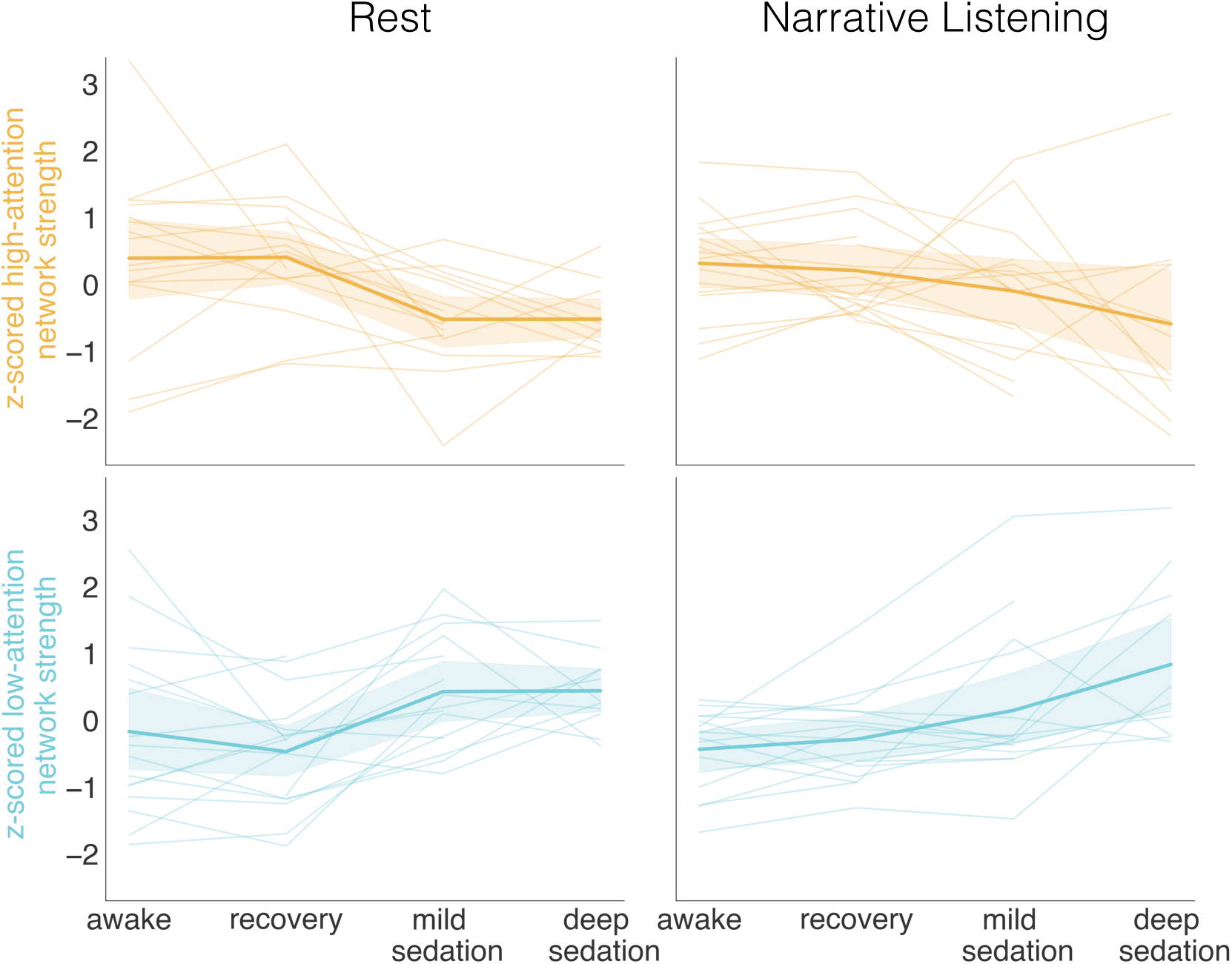
High-attention and low-attention network strength during two tasks (rest, narrative listening), and four sedation conditions (*awake, recovery, mild sedation*, and *deep sedation*). Network strength values were *z*-scored within graph for visualization. Semi-transparent lines represent individual participants, Bold lines represent average across participants, with the shaded region indicating the 95% confidence interval. Some semi-transparent lines do not extend the full length of the plot as not all participants had complete data for all sedation and task conditions.

To account for potential effects of head motion on functional network strength, we performed two control analyses. First, we regressed mean framewise displacement and fraction of censored frames from high- and low-attention network strength scores. We then repeated the analysis described above with the residuals of this regression, performing paired *t*-tests comparing residualized attention network strength while individuals were awake and under deep sedation during rest and narrative listening. Demonstrating that effects of propofol on attention network strength are robust to effects of head motion, residualized high-attention network strength during rest was higher during the *awake* than the *deep sedation* condition (rest: *t*_9_ = 3.56, *p* = 0.006; narrative: *t*_9_ = 2.73, *p* = 0.023). We observed the opposite pattern of results for residualized low-attention network strength (rest: *t*_9_ = -2.34, *p* = 0.044; narrative: *t*_9_ = -3.59, *p* = 0.006).

As a second motion control, we replicated the mixed effects analysis including mean framewise displacement and fraction of censored frames in each run as predictors. Results were consistent with those described above: models revealed a main effect of sedation condition wherein less sedation was associated with functional connectivity signatures of stronger attention (high-attention: *b* = -0.82, SE = 0.16, F[3,93.7] = 9.87, *p* = 1.02×10^−5^; low-attention: *b* = 0.68, SE = 0.15, F[3,92.9] = 8.5, *p* = 4.73×10^−5^). There was no main effect of task or interaction of sedation condition and task on high-attention network strength. For low-attention network strength, there was a main effect of task (*b* = -0.15, SE = 0.07, F[1,92.3] = 4.1, *p* = 0.045) such that low-attention network strength was higher during rest than narrative-listening scans.

### Propofol specifically modulates functional connectivity signatures of sustained attention

#### Propofol modulates sustained attention networks more than fluid intelligence networks

Is propofol’s effect on the connectome *specific* to networks predicting sustained attention? We tested this question in three ways. First, we investigated propofol’s effect on a functional connectivity network defined in previous work to predict fluid intelligence (Greene et al., 2018). Attention function varies across individuals but also varies within a single individual over time, and the sustained attention CPM is sensitive to both these individual differences and intraindividual variability (Rosenberg et al., 2020). Fluid intelligence, by comparison, may reflect a more trait-like aspect of behavior, as evidence suggests it is relatively stable within an individual over time (Kazlauskaite & Lynn, 2002; Schaie et al., 2004). Consequently, drug induced changes in cognitive state may modulate the fluid intelligence networks to a lesser degree than they do the sustained attention networks.

In both the resting-state and narrative-listening conditions, high-fluid intelligence network strength did not significantly differ between *awake* and *deep sedation* scans (rest: *t*_9_ = .27, *p =* 0.79, narrative listening: *t*_9_ =.61, *p =*.56) (**Fig. 2**). Low-fluid intelligence network strength was significantly greater in the *deep sedation* scans for the narrative-listening condition only (rest: *t*_9_ = -1.78, *p =* 0.11, narrative listening: *t*_9_ = -2.58, *p =* 0.0297). A mixed effect models of propofol’s effect on fluid intelligence network strength did not show a main effect of sedation condition for high-fluid intelligence network strength, but did show a main effect of sedation for low-fluid intelligence network strength, wherein greater sedation was associated with higher low-fluid intelligence network strength scores (high-fluid intelligence: *b* = - 0.19, SE = 0.18, F[3, 94.5] = .44, *p* = 0.72; low-fluid intelligence: *b* = 0.56, SE = 0.18, F[3, 92.4] = 4.52, *p* = 0.005). There was no main effect of task or interaction of sedation condition and task on high-fluid intelligence or low-fluid intelligence network strength.

**Figure 2.**
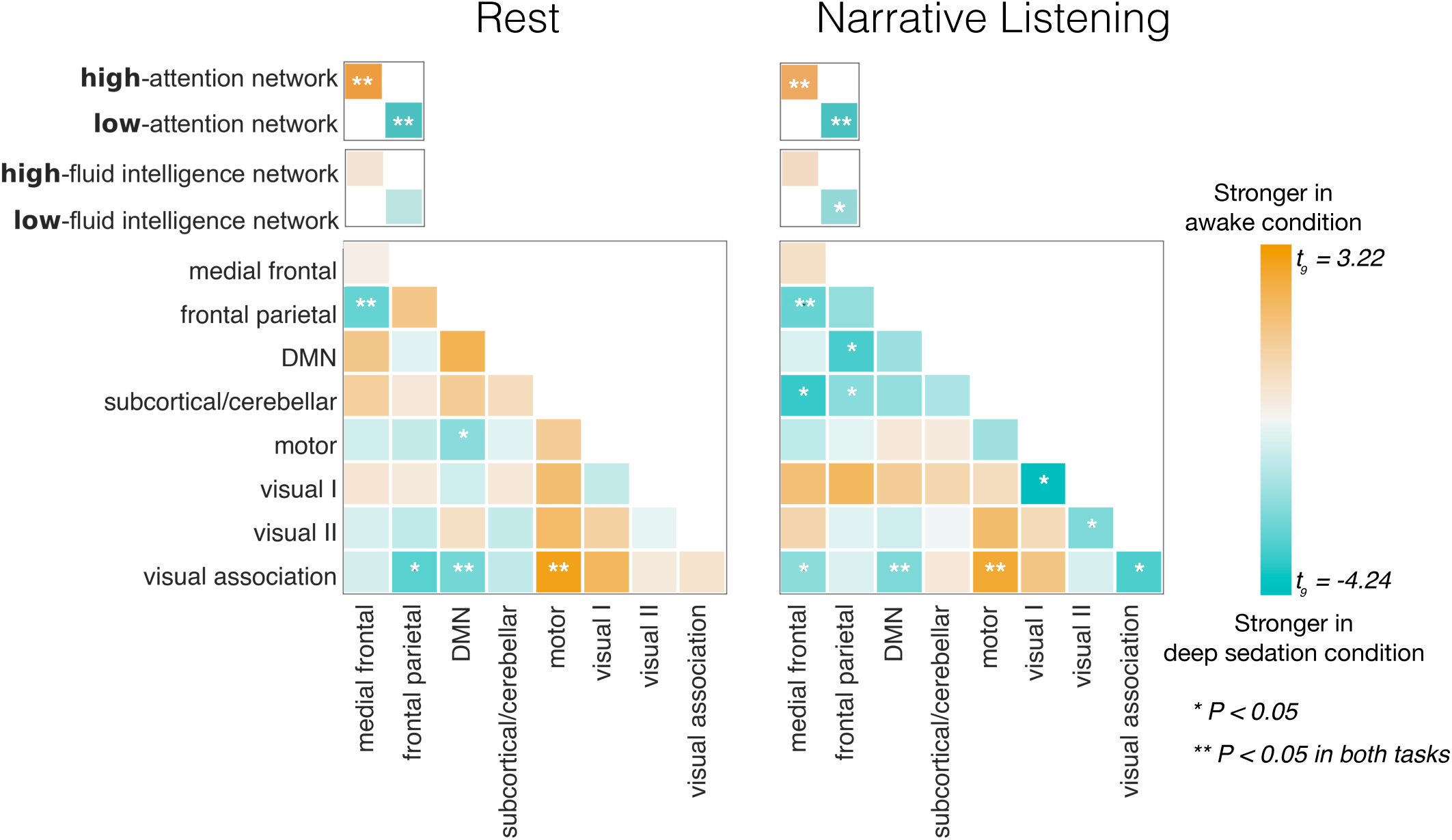
Effects of propofol on functional network strength. Differences in within-network and between-network strength (averaged functional connectivity) during the *awake* and *deep-sedation conditions*.

We performed paired *t*-tests comparing the percent change in overall sustained attention and fluid intelligence network strength from *awake* to *deep sedation* during both task conditions. Overall network strength was calculated as the difference between strength in the network predicting higher behavioral scores and the network predicting lower behavioral scores. Results revealed greater percent change in sustained attention than fluid intelligence network strength during both rest and narrative listening (rest: *t*_9_ = 2.27, *p =* 0.049, narrative listening: *t*_9_ =3.61, *p =*.006). Thus, propofol has a greater effect on the sustained attention networks than networks predicting fluid intelligence. (We did not test for a 3-way interaction between functional network, task, and sedation condition with mixed effects models as only 17 participants were included in this analysis.)

#### Propofol modulates sustained attention networks more than canonical resting-state networks

Although networks defined to predict fluid intelligence were not as sensitive to sedation as the sustained attention networks, this does not preclude the possibility that other functional networks are affected by propofol sedation—perhaps even more so than the sustained attention networks. To test this possibility, we assessed the effect of sedation on functional connectivity in eight canonical resting-state networks as well as the pairwise connections between these networks (36 network pairs in total) (**Fig 2**). Providing further evidence for the specificity of the effect on the sustained attention CPM, propofol’s effect on the high-attention network was numerically greater than the positive effects for any of the 36 pairs of canonical resting-state networks during both tasks. The effect of propofol on the low-attention network was greater than any negative effect for all pairs of canonical resting-state networks during rest, and all but one network during narrative listening (within primary visual network connections).

Finally, we compared differences in sustained attention network strength between the *awake* and *deep sedation* conditions (assessed with a paired *t*-test) to differences in the strength of in 10,000 same-size random networks. During rest, the effect in the high-attention network was greater than 96.71% of same-sized random networks, and the effect in the low-attention network was greater than 98.79% of same-sized random networks. During narrative listening, the effect in the high-attention network was greater than 99.79% of same-sized random networks, and the effect in the low-attention network was greater than 100% of same-sized random networks. In sum, this replication demonstrates the robustness of the finding that the sustained attention CPM is uniquely sensitive to propofol sedation.

Beyond the sustained attention networks, effects of propofol on functional connectivity on canonical networks are consistent with those reported previously (Rosenberg et al., 2020). This prior work, performed in an independent dataset, found that seven of the 36 canonical network pairs were significantly modulated by propofol administration. Of those seven pairs, six overlap with the seven network pairs showing the greatest effect in the present study during rest (*p* < 0.07, 1. frontoparietal-visual association, 2. motor-visual association, 3. frontoparietal-medial frontal, 4. default-visual association, 5. motor-default, 6. default-default, 7. visual I-visual association). None of the effects in the canonical networks in the present study survived Bonferroni correction for 36 tests (**Fig. 2**).

#### Propofol sedation networks overlap with the sustained attention networks

The sustained attention networks are modulated by propofol sedation, but to what extent do the functional connections that change most with propofol sedation overlap with the connections in the sustained attention CPM’s high- and low-attention networks? To investigate, we examined the overlap between both the *Awake* networks (i.e., the network stronger during wakefulness) and the *Deep Sedation* networks (i.e., the network stronger during deep sedation) and the high- and low-attention networks. We predicted that the both the high-attention network would share significant overlap with the *Awake* networks, while the low-attention network would share significant overlap with the *Deep Sedation* networks.

To this end, we used the network-based statistic to identify functional networks most affected by propofol sedation. We did this at three different edge-selection thresholds, retaining edges that significantly differed between *Awake* and *Deep Sedation* scans at *p* = .05, .01, .001 respectively. We detected significant *Awake* networks (network significance defined as *p* < .05) for all three edge-selection threshold levels for the rest condition, and one out of three (.01) edge-selection threshold levels for the narrative-listening condition (**Figs. 3, 4**). We detected significant *Deep Sedation* networks for all three edge-selection threshold levels tested and for both task conditions. In general, *Awake* networks consisted of nodes in the occipital, prefrontal, and temporal lobes, while *Deep Sedation* networks, while *Deep Sedation* networks were dominated by nodes in the prefrontal, limbic, and temporal lobes.

**Figure 3.**
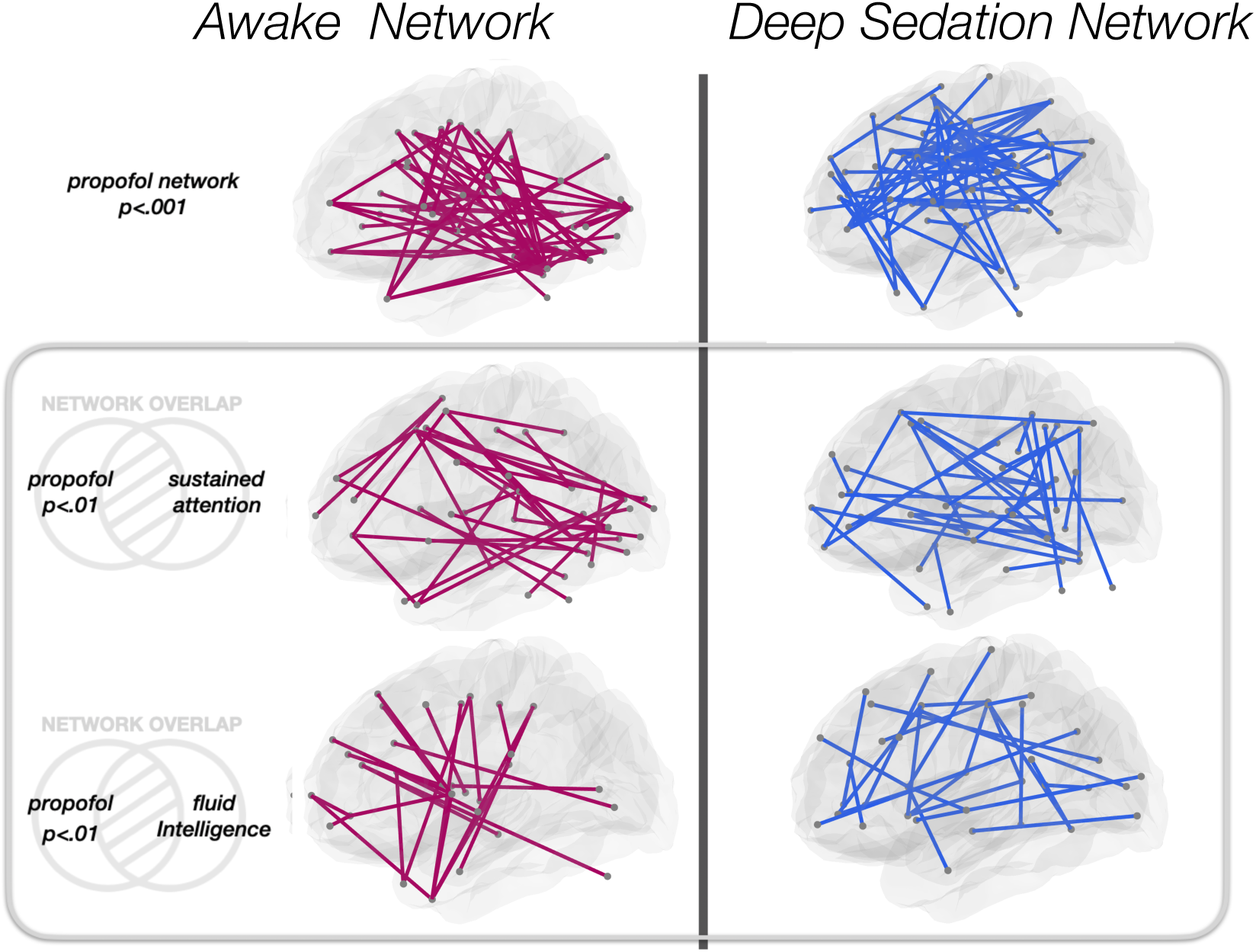
The *Awake* network shows significant overlap with the low-attention and low-fluid intelligence networks, whereas the *Deep Sedation* network shows significant overlap with the low-attention and low-fluid intelligence networks.

**Figure 4.**
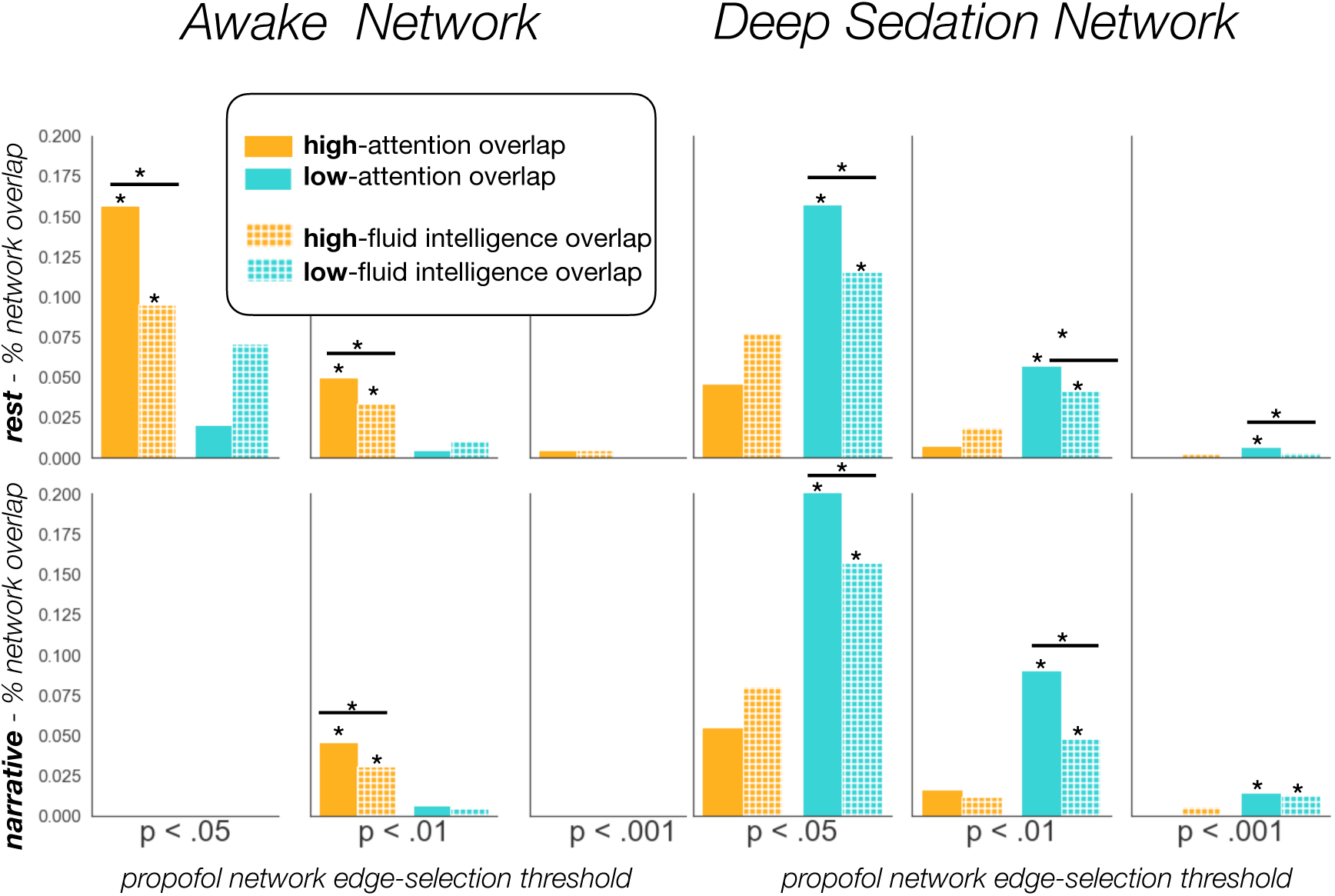
The sustained attention networks show significantly greater overlap with the propofol networks than do the fluid intelligence network. P-values in the x-axis indicate the threshold at which edges were selected when defining the propofol networks. Stars indicate the significance of the percent overlap (*p* < .05, Bonferroni corrected for 4 comparisons, *p*-values computed using hypergeometric cumulative density function). Stars above black lines indicate a significant difference in percent overlap between networks (*p* < .05, p-values computed using permutation testing).

As expected, there was significant overlap between *Awake* networks and the high-attention network but not the low-attention network (**Fig. 4**). Furthermore, there was significant overlap between *Deep Sedation* networks and the low-attention network but not the high-attention network.

For the .01 threshold *Awake* rest network, 37 edges overlapped with the high-attention network, making up 4.9% of the high-attention network (*p* = 1.25 × 10^−7^). The reverse pattern held true for the *Deep Sedation* network, with 36 edges in common between the .01 threshold *Deep Sedation* network and the low-attention network, making up 5.7% of the low-attention network (*p* = 6.45 × 10^−9^). This result replicated in the narrative listening condition (high-attention *Awake* overlap: 34 edges, 4.5% of network, *p* = 1.11 × 10^−9^; low-attention *Deep Sedation* overlap: 57 edges, 9.0% of network, *p* = 9.44 × 10^−15^). Consequently, the functional networks that differ between sedation and wakefulness overlap with those that differ between states of better and worse sustained attention.

We expected the propofol networks to show less overlap with the fluid intelligence networks than the sustained attention networks, in line with the hypothesis that the sustained attention networks are more sensitive to cognitive and attentional state changes. We observed significant overlap between *Awake* networks and the high-fluid intelligence network but not the low-fluid intelligence network. For the .01 threshold *Awake* rest network, there were 23 edges overlapping with the high-fluid intelligence network, making up 3.4% of the high-fluid intelligence network (*p* = 0.007). The *Deep Sedation* network presented the opposite pattern, showing significant overlap with the low-fluid intelligence but not the high-fluid intelligence network. For the .01 threshold *Deep Sedation* rest network, there were 27 edges shared with the low-fluid intelligence network, making up 4.2% of the low-fluid intelligence network (*p* = 9.4 × 10^−5^). These patterns replicated in the narrative listening condition (high-fluid intelligence *Awake* overlap: 21 edges, 3.1% of network, *p* = 2.37 × 10^−4^; low-fluid intelligence *Deep Sedation* overlap: 31 edges, 4.8% of network, *p* = 9.44 × 10^−15^).

Supporting our prediction, comparisons of network overlap revealed that the sustained attention network showed significantly greater overlap with the propofol networks in 8 the 10 comparisons (2 propofol networks [*Awake, Deep Sedation*] x 3 edge selection thresholds [*p* < .05, .01, .001] x 2 task conditions [rest, narrative listening], minus two instances where no significant propofol network was found = 10 comparisons), and numerically greater overlap for all comparisons except overlap with the .001 threshold *Awake* rest network. This difference is not merely driven by a difference in the overall size of the sustained attention and fluid intelligence networks: the networks are comparable sizes (sustained attention networks = 1387 edges, fluid intelligence networks = 1333 edges), and this difference persists when comparing overlap as a percentage of network size. These results suggest that the functional networks most strongly modulated by propofol share a more similar architecture to the functional networks underlying sustained attention function than to those associated with fluid intelligence.

### Propofol and task condition modulate between-subject functional connectome similarity

In addition to gauging propofol’s effect on neural signatures of sustained attention, we assessed how sedation affects functional connectivity patterns more broadly. To this end, we examined the between-subject connectome similarity while participants were awake versus sedated. A paired *t*-test comparing similarity values during the *awake* resting-state scans to those of *deep sedation* resting-state scans demonstrated that participants are more similar to one another when awake then when under sedation (*t*_9_ = 14.96, *p* = 1.15×10^−7^). Furthermore, this result replicated in the narrative-listening condition (*t*_9_ = 6.43, *p* = 1.21×10^−4^). Interestingly, this result suggests that propofol sedation does not simply mute individual differences in functional connectivity patterns and cause all participants to show a connectivity pattern common across the population. Instead, participants’ functional connectivity patterns look more similar to each other when they are awake.

In addition to assessing the effect of sedation on similarity, we also examined the effect of task. The paired *t*-test revealed, surprisingly, greater similarity during resting-state scans than during narrative-listening scans, and this result replicated in three of the four sedation conditions (awake: *t*_15_= 6.89, *p* = 5.17×10^−6^; mild: *t*_12_= .50, *p* = .62; deep: *t*_7_= 6.64, *p* = 2.935×10^−4^; recovery: *t*_14_= 9.84, *p* = 1.13×10^−7^). This finding is somewhat counterintuitive given extensive work showing that individuals shown the same movie or played the same audio narrative in the scanner demonstrate higher levels of inter-subject correlation compared to rest (Hasson et al., 2004; Kandeepan et al., 2020). In these data, Kandeepan et al. (2020) observed significant inter-subject correlation at all levels of sedation, but not during rest. Furthermore, it has been demonstrated previously in a separate dataset that functional connectome similarity is increased both within and across participants during movie-watching compared to rest (Vanderwal et al., 2017). This analysis suggests that greater BOLD time course similarity may not necessarily translate to greater functional connectome similarity between individuals.

## Discussion

To what extent does an individual’s functional connectome reflect stable individual traits vs. varying mental states? Work demonstrates that individual differences can be discerned from functional connectivity patterns, with successful prediction of between-subject variation in personality (Cai et al., 2020; Hsu et al., 2018), working memory performance (Avery et al., 2019; Galeano Weber et al., 2017; Yamashita et al., 2018) and reading recall (Jangraw et al., 2018), among other behaviors. But what about intra-individual differences? Here we add support for the existence of state-specific variation in functional connectome by testing if a predefined network predicting sustained attention is sensitive to pharmacological intervention with propofol. Using an independent dataset, we replicate the effect of propofol on the sustained attention CPM. We find, as predicted in our preregistration, that greater propofol sedation is associated with neural signatures of worse sustained attention function, suggesting that behaviorally relevant aspects of the functional connectome vary with short-term, pharmacologically induced changes in cognitive and attentional states.

The utility of a connectome-based model of cognitive function relies on it being both generalizable and specific. Generalizing across datasets, for instance, provides evidence that a model characterizes the measure of interest, rather than merely picking up on idiosyncrasies in a given participant sample from a single fMRI scanner. In the present study, we demonstrate cross-dataset generalizability by replicating the effects of propofol on the sustained attention networks in data from a novel set of participants collected at an independent scanning site. Just as cross-dataset generalizability is important, cross-measure generalizability is also key. A model is less useful if it only predicts a specific behavioral measure (such as psychological task performance), rather than other measures of the same underlying cognitive or attentional function (such as ADHD symptomatology for models of sustained attention) (Rosenberg, Finn, et al., 2016). An emphasis on generalizability both across datasets and related measures, among other practices such as preregistration, is crucial for tackling the reproducibility crisis in psychology and neuroscience.

Though perhaps less frequently discussed, the specificity of a model is as important as its generalizability. If a model generalizes in a way such that it cannot be distinguished from other models (here, predefined predictive networks), it is also less useful if one aims to disentangle various cognitive functions from one another. For instance, suppose one wishes to characterize the functional networks associated with mathematical ability. If a predictive model performs equally well in predicting individual’s math and reading scores, it is hard to argue that this model informs our understanding of the neural underpinnings of mathematical aptitude per se. Critically, in the current work, we also add evidence to the “specificity” criteria, as we demonstrate that not all behaviorally predictive brain networks are equally sensitive to propofol-induced state changes. Rather, our results suggest that canonical resting-state networks as well as a previously defined model of fluid intelligence are less sensitive than the sustained attention network to the effects of propofol sedation. This suggests that perhaps the model of fluid intelligence captures less state-like variability and more trait-like variability compared to the sustained attention CPM, in line with research demonstrating that fluid intelligence is relatively stable within individuals’ over time (Kazlauskaite & Lynn, 2002; Schaie et al., 2004). Furthermore, after defining “propofol” networks, based on the networks differing most between deep and awake conditions, we found that the propofol network had greater overlap, on average, with the sustained attention networks than the fluid intelligence networks. These results suggest that the functional networks underlying attention function, compared to those associated with fluid intelligence, may share a more similar architecture to the functional networks most strongly modulated by sedation with propofol.

Future work will reveal the extent to which networks defined to predict individual differences in other processes are sensitive to within-subject state change. Although a growing body of work has found evidence that the connectome-based model of individual differences in sustained attention function also successfully differentiates between within-subject state change, this may not be the case for all behavioral traits. For instance, 3,4-Methylenedioxymethamphetamine (MDMA) influences social function, increasing subjective sociability and feelings of closeness with others (Bedi et al., 2009). Would a functional network defined to predict trait extraversion (Hsu et al., 2018) be sensitive to within subject state change induced by administration of MDMA? Perhaps the neural basis of individual differences in social function are orthogonal to the neural basis of change in social behavior within one person. More broadly, pharmacological studies provide unique opportunity to test the generalizability and specificity of predictive models.

The current study differs somewhat with respect to prior findings regarding sedation’s effect on functional connectivity patterns. For instance, past work found a decrease in within-default-mode-network (DMN) connectivity with propofol sedation (Boveroux et al., 2010; Guldenmund et al., 2017; Qiu et al., 2017; Tang & Ramani, 2016). However, results here suggest DMN connectivity with other networks is differentially affected by propofol sedation. Specifically, within-DMN connectivity decreased during rest but not narrative-listening scans, whereas DMN-visual association network connectivity increased during both tasks. This difference in findings can perhaps be attributed the fact that lower levels of sedation have been associated with no change in DMN connectivity (Stamatakis et al., 2010). The present study defines “deep” sedation as a lower level of sedation (Ramsay level 5) compared to the sedation condition in some prior work (Ramsay level 5-6 in Boveroux et al., for instance, or “deep sedation” as defined by the American Society of Anesthesiologists in Qiu et al., 2017) reporting decreases in DMN connectivity. Results in the present study are also contrast with prior work showing that sedation decreases frontoparietal connectivity (Amico et al., 2014; Boveroux et al., 2010). Data here suggest increased frontoparietal connectivity during sedation, specifically frontoparietal-medial frontal connectivity, as well as frontoparietal-visual association connectivity. Observed changes in functional connectivity in visual regions, however, are consistent with past work, where sedation was associated either with no change, or an increase in connectivity (Martuzzi et al., 2010; Qiu et al., 2017). Here we report significant increases in within-network connectivity in V1, V2, and visual association networks in the narrative listening condition, and no significant changes in visual network connectivity during rest. Additional work may resolve how parameters such as sedation level, scan length, preprocessing approach, and task mediate the effect of propofol sedation on functional connectivity. While the present findings are not wholly consistent with all prior work, propofol’s effect on functional connectivity patterns is consistent across tasks within the current dataset (rest and narrative listening) and with the findings reported in an independent dataset analyzed with a similar preprocessing pipeline and data exclusion criteria (see *Propofol modulates sustained attention network to a greater degree than canonical resting state networks*) (Rosenberg et al., 2020).

In addition to investigating the effect of propofol on connectome-based models of behavior and canonical resting-state networks, we characterized how sedation and task manipulation affected the similarity of individuals’ overall functional connectivity patterns. With regards to the effect of sedation on similarity, we began with two plausible competing hypotheses: 1) Sedation decreases ongoing thoughts, feelings, etc., that may differ across individuals and drive differences in functional connectivity patterns, thereby resulting in more similar connectivity patterns across individuals. 2) Any kind of cognitive processing (even if the content differs to across individuals) acts as a constraint on functional connectivity, resulting in connectome patterns that are more similar in the absence of sedation. We found evidence for the latter hypothesis, observing greater similarity when participants were awake compared to when they were sedated. In addition to examining the effect of task on between-subject similarity, our analyses explored the extent to which task condition affects between-subject similarity. We found that rest was associated with greater across-subject connectome similarity compared to narrative-listening. This finding is surprising, given past work that demonstrates that, in development, movie-watching raises between-subject connectome similarity relative to rest (Vanderwal et al., 2017). Lack of visual input in the present study may help explain this difference in results. In Vanderwal et al., 2017, participants watched and listened to a film, whereas here, participants only listened to audio for the “narrative” condition. Synchronous visual input may drive between-subject similarity higher in the “narrative” condition relative to rest. Furthermore, in Vanderwal et al., 2017, the resting-state scan was collected while participants stared at a fixation cross, while in the present study participants were instructed to keep their eyes closed. Past work has found significant differences between functional connectivity for eyes open vs. eyes closed resting-state scans, and perhaps this helps explains the divergence of the present findings from prior work (Costumero et al., 2020). One limitation of the present study is a relatively small sample size (*n* = 17). Thus, additional work can help determine the ways in which tasks and pharmacological interventions modulate connectome similarity between participants.

In sum, we replicate the effect of propofol on the sustained attention CPM and provide evidence that the magnitude of this effect is specific to the sustained attention network. Additionally, we demonstrate that task and sedation manipulations in this dataset have significant and unexpected effects on connectome similarity. Ultimately, we add greater support for the idea that within-subject attentional state changes are encoded in within-subject differences in functional connectivity and offer grounds for further exploration of the way functional connectivity models of different cognitive functions may differ in their sensitivity to state-vs. trait-like differences.

## Data Availability

Code will be made available on GitHub upon publication: https://github.com/tchamberlain/propofol. MRI data are available on OpenNeuro: https://openneuro.org/datasets/ds003171/

## Acknowledgements

This work was supported by National Science Foundation BCS-2043740 (M.D.R.) and resources provided by the University of Chicago Research Computing Center. We thank Abigail Greene for sharing the connectome-based predictive models of fluid intelligence.

## Supplementary Material

**Supplementary Figure 1.**
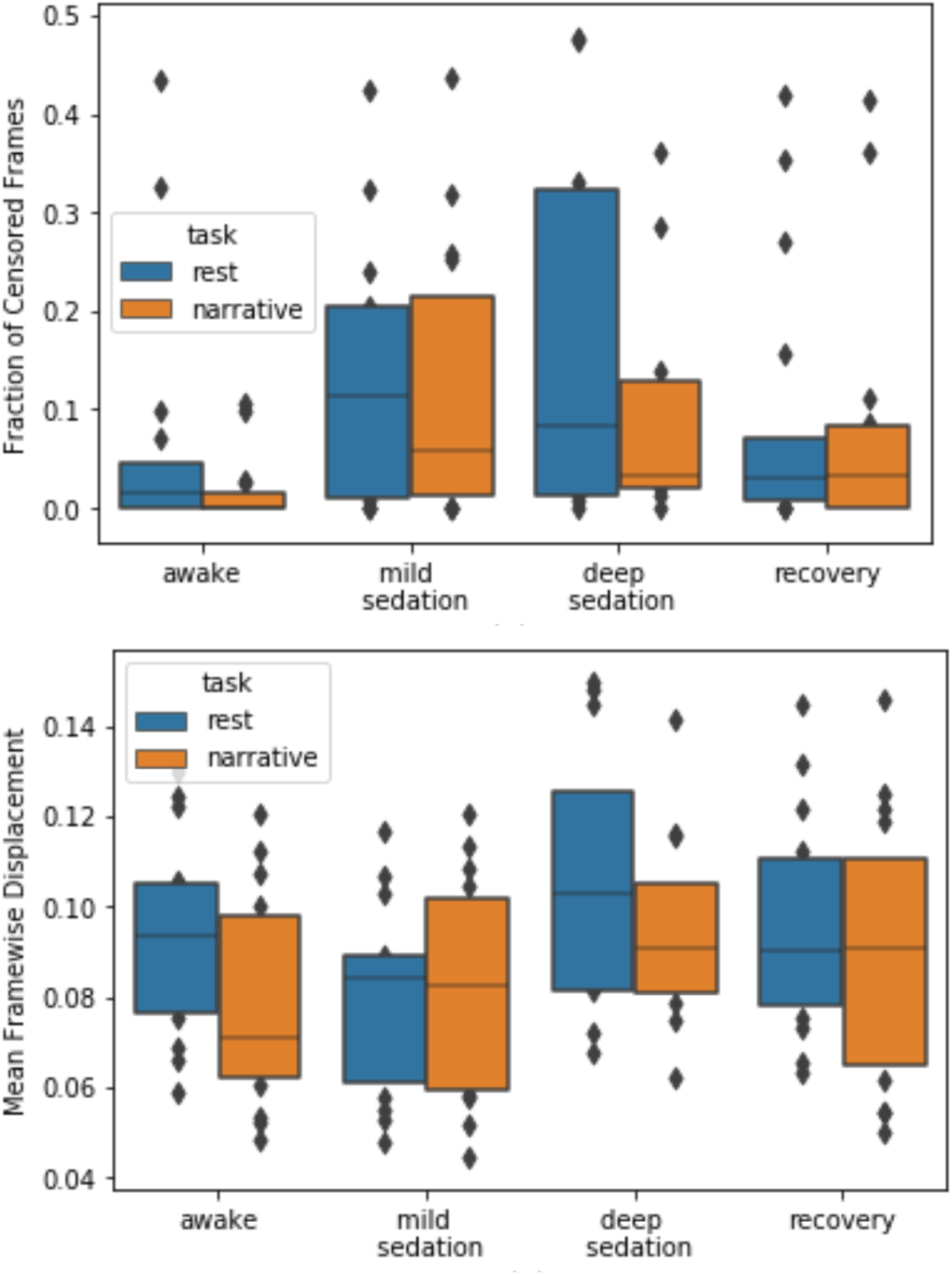
Head motion across sedation conditions (mean framewise displacement, and fraction of frames censored). See methods for frame-censoring criteria.

**Supplementary Figure 2.**
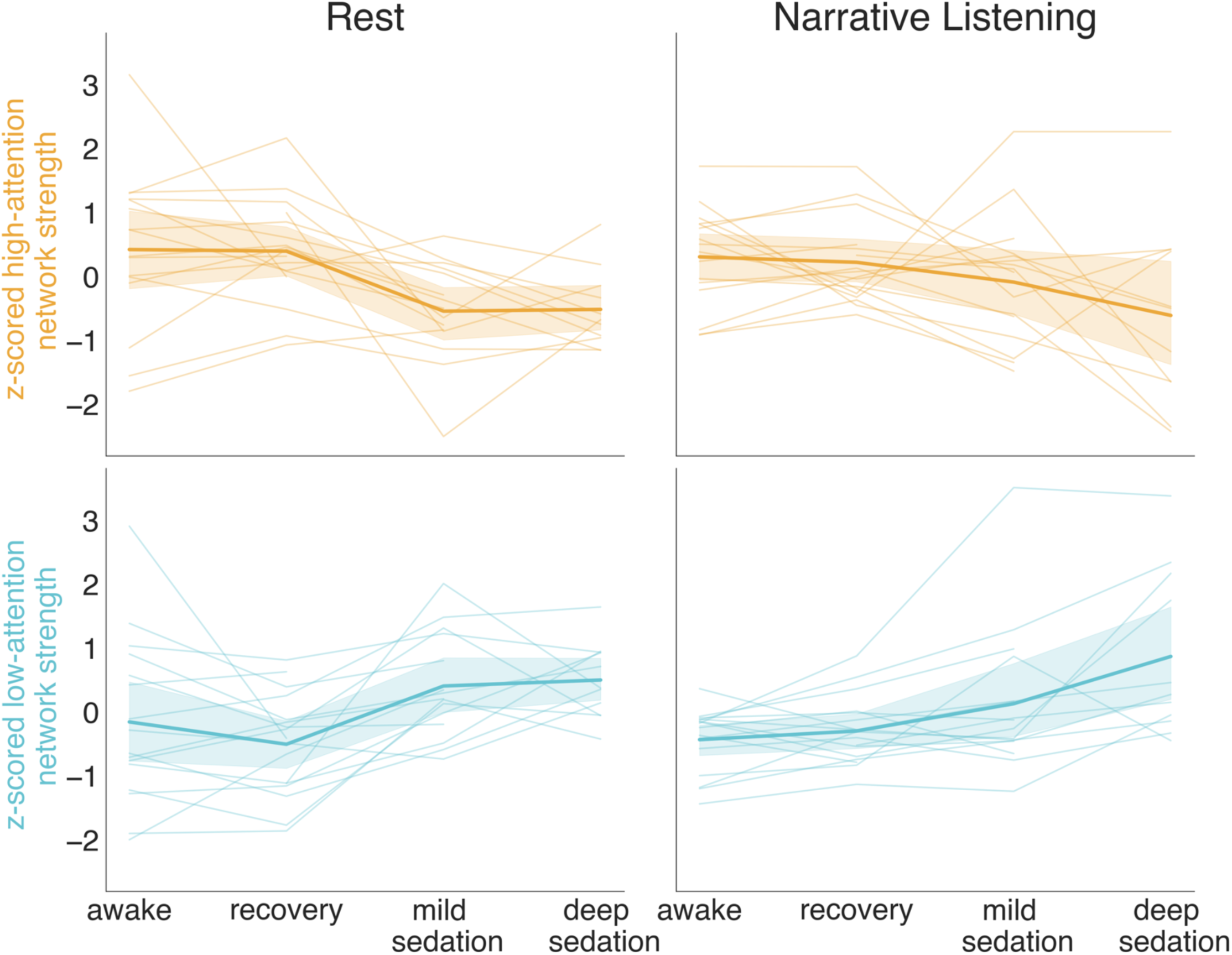
Effect of propofol on network strength: excluding all missing nodes. We repeated our primary replication analysis after dropping any nodes in the 268-node Shen parcellation which were missing in any scan. 30 nodes were missing in total, leaving a 238 × 238 functional connectivity matrix from which we calculated network strength scores (see methods). Results were consistent with those obtained using all 268 nodes. The high-attention network strength during rest was greater during the awake than the deep sedation condition (*t*_9_ = 3.22, *p =* 0.010). The low-attention network exhibited the opposite pattern of results (*t*_9_ = -3.98, *p* = 0.003). This pattern of results replicated in the narrative-listening condition (high-attention: *t*_9_ = 2.62, *p* = 0.028; low-attention: *t*_9_ = -4.01, *p* = 0.003).

